# Structural Evolution of LEAFY Reveals DNA-Mediated Cooperativity and Dimerization Shifts at the Water-to-Land Transition

**DOI:** 10.64898/2026.04.13.717636

**Authors:** Leonie Verhage, Emmanuel Thévenon, Hicham Chahtane, Loïc Grandvuillemin, Max H. Nanao, Renaud Dumas, Chloe Zubieta, François Parcy

**Affiliations:** Laboratoire Physiologie Cellulaire et Végétale, Univ. Grenoble Alpes, CNRS, CEA, INRAE, IRIG-DBSCI-LPCV, 17 avenue des martyrs, 38054, Grenoble, France; Green Mission Pierre Fabre, Conservatoire Botanique Pierre Fabre, Institut de Recherche Pierre Fabre, Soual, France; IBS, Univ. Grenoble Alpes, CNRS, CEA, Grenoble, France; Structural Biology Group, European Synchrotron Radiation Facility, 71 Avenue des Martyrs, 38000 Grenoble, France

## Abstract

The evolution of transcription factor (TF) DNA-binding specificity is a major driver of gene regulatory innovation. Unlike most TFs, which diversify through gene duplication and neofunctionalization, the plant-specific LEAFY (LFY) TF evolved novel binding specificities without extensive duplication. Here, we combine experimental structural determination and biochemical assays to reveal how LFY’s dimerization and DNA-binding preferences shifted during the water-to-land transition. We present crystal structures of the LFY DNA-binding domain (DBD) from the hornwort *Nothoceros aenigmaticus* and the alga *Interfilum paradoxum* bound to DNA, demonstrating two distinct dimerization mechanisms: one mediated by direct protein-protein interactions and another driven by DNA-mediated cooperativity. In the ancestral state, LFY likely bound DNA as a dimer through DNA-mediated cooperativity, with protein-protein dimerization emerging later, enforcing new DNA-binding preferences. Our findings support a revised evolutionary scenario for LFY, highlighting the dynamic interplay between protein-DNA and protein-protein interactions as key drivers of TF binding specificity. This work deepens our understanding of how structural adaptations in TFs underpin evolutionary transitions in gene regulation.

## Introduction

Innovation in gene expression is a fundamental driver of evolutionary adaptation across living organisms. Transcription factors (TFs) are key actors in this process: these proteins regulate gene expression by binding to specific DNA sequences—known as *cis*-elements or TF binding sites (TFBS)— most often present in the promoter regions of the genes they regulate. Many TFs act as dimers (homo- or heterodimers) or higher order oligomers (Blanc-Mathieu et al. 2024; Xie et al. 2025). Consequently, their DNA binding site preferences change due to the presence of multiple sterically constrained DNA-binding domains (DBD) leading to an increased specificity that reads out both DNA sequence and spatial organization of the multiple TFBS. Evolutionary changes in gene expression can arise from modifications in *cis*-regulatory elements, alterations in the DNA-binding specificity of TFs themselves or both (reviewed in (Hill et al. 2021)). Many TFs tend to retain the same DNA binding specificity over large evolutionary times (Nitta et al. 2015; Zenker et al. 2024). However, transcription factors can also undergo functional diversification through changes in their DNA-binding residues or alterations in their oligomerization and protein-protein interaction properties (Nakagawa et al. 2013; Gera et al. 2022). For example, even subtle amino acid substitutions in the DNA-binding domain can dramatically shift binding specificity. Nakagawa et al. (2013) demonstrated in forkhead transcription factors that evolutionary changes in key residues led to the recognition of novel cis-regulatory elements. Similarly, Gera et al. (2022) showed that modifications in oligomerization interfaces—such as those observed in whole-genome duplicated transcription factors—can rewire regulatory networks by enabling new protein-protein interactions or altering the stability of TF complexes. These studies illustrate how structural adaptations in TFs can drive the evolution of gene regulatory networks and contribute to phenotypic novelty.

A compelling example of a TF that experienced several types of changes in DNA- and protein-protein interaction specificity is the plant-specific TF, LEAFY (LFY). LFY is present in all land plants and originated in streptophyte algae (Leebens-Mack et al. 2019; Lai et al. 2020; Baumgart et al. 2024). But unlike most TFs, LFY did not expand into a multigene family in land plants but instead remained low-copy across most species. While its functional role in algae remains elusive, LFY plays critical roles in land plants: it regulates cell division in the moss *Physcomitrium patens* and apical growth in the fern *Ceratopteris richardii* (Tanahashi et al. 2005; Plackett et al. 2018; McConnell et al. 2026). In angiosperms, LFY is essential for floral meristem identity and flower development (Denay et al. 2017; Rieu et al. 2023). Its function relies on two conserved domains: an N-terminal Sterile Alpha Motif (SAM) domain, which mediates oligomerization, and a C-terminal Helix-Turn-Helix DNA-binding domain (DBD) that dimerizes upon DNA binding (Hamès et al. 2008; Sayou et al. 2016). The SAM domain is tethered to the DBD via a flexible linker. While the SAM domain favors LFY binding to clustered sites, it does not affect DNA-binding specificity (Sayou et al. 2016). The DBD and its dimerization mode, in contrast, are central to LFY’s ability to bind specific DNA sequences and to regulate target genes, yet how its binding specificity evolved remains poorly understood. Despite the high conservation of its DBD, LFY has undergone remarkable shifts in DNA-binding specificity over evolutionary time (Sayou et al. 2014; Brunkard et al. 2015; Gao et al. 2019). This is attested by the diverse binding motifs recognized by LFY across plant lineages: in liverworts and vascular plants such as *Arabidopsis thaliana*, LFY binds, as a dimer, a 19-bp palindromic type I site with a 3-bp spacer separating inverted half-sites (Fig. 1A). In *Physcomitrium patens*, the PpLFY1 dimer (thereafter called PpLFY) recognizes a palindromic type II site, also with a 3-bp spacer but featuring a distinct half-site sequence. In the alga *Klebsormidium subtile*, KsLFY-DBD dimer binds a type III motif—a palindromic site lacking the 3-bp spacer. Notably, the hornwort *Nothoceros aenigmaticus* NaLFY exhibits a promiscuous binding, defined as the ability to form dimeric complexes on all three motif types (Sayou et al. 2014). Structural studies have elucidated how LFY binds to type I (for *Arabidopsis thaliana* AtLFY, PDB 2VY1) and type II (for *Physcomitrium patens* PpLFY, PDB 4BHK) motifs, revealing a conserved dimer configuration but distinct protein-DNA contacts (Sayou et al. 2014). However, the promiscuous binding of *Nothoceros aenigmaticus* NaLFY as well as the DNA binding of algal LFY that exhibit unique residues at DNA-contacting positions remain structurally uncharacterized. Moreover, the absence of a 3-bp spacer in the type III motif implies a distinct spatial arrangement of the two DNA-binding domains (DBDs), raising questions about whether these DBDs interact through a novel dimerization interface.

**Figure 1.**
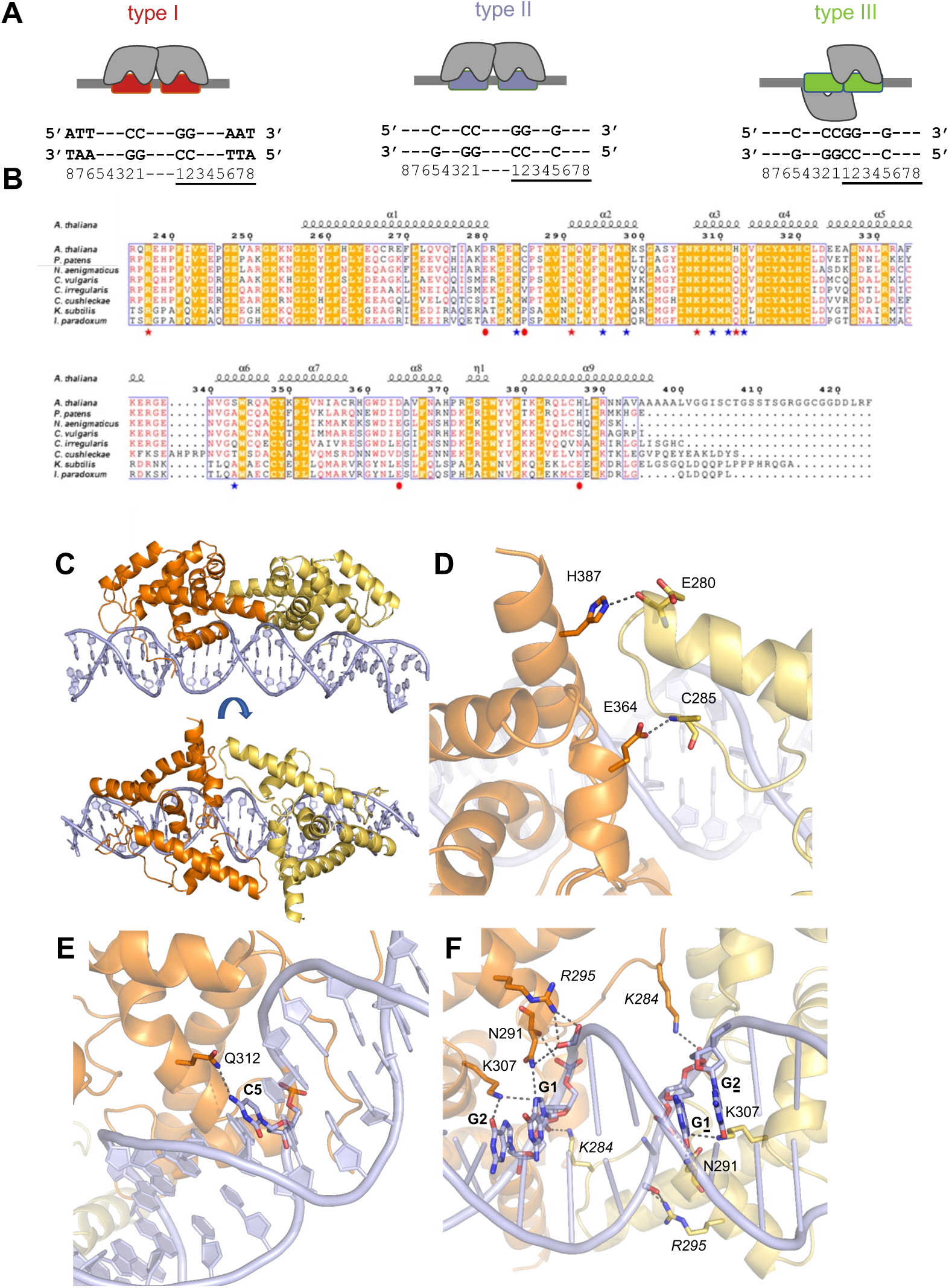
Structural basis of NaLFY-DBD binding to type II DNA motif. (A) Schematic of different binding types with the LFY-DBD depicted in grey and the motifs in different colors: type I in red, type II in blue and type III in green (this color code will be used throughout the manuscript). Bases are numbered for each half-site (underlined or not). (B) Alignment of LFY-DBD protein sequences from land plants (*A. thaliana* as flowering plant, *P. patens and N. aenigmaticus* as bryophytes and others as charophyte algae). The plain circles indicate residues involved in DBD homodimerization in type I and II motifs, red stars indicate residues interacting with DNA bases and blue stars, residue interacting with the phosphate backbone. (C) Two views of the crystal structure of NaLFY-DBD (yellow and orange) bound to type II DNA (blue). (D) Close-up on the residues involved in the dimerization interface. The side chain of Glutamate 364 (E364) and Histidine 387 (H387) interacts with the main chain of Cysteine 285 (C285) and Glutamate 280 (E280), respectively. (E) Close-up on specificity determining residues (Glutamine, Q312) and DNA base (desoxycytosine, C5). (F) Close-up on residues interacting with DNA in the major groove. Plain residues interact with the bases and italicized ones with the phosphate backbone.

Based on available data, an evolutionary trajectory was initially proposed in which LFY ancestrally bound the type III motif as a dimer, transitioning to types I and II via a promiscuous intermediate capable of recognizing all three motifs. This scenario has been revisited following the discovery of additional *LFY* genes that indicate a LFY duplication event around the water-to-land transition (Brunkard et al. 2015; Gao et al. 2019). However, several critical steps in LFY’s evolutionary history remain unresolved.

To address central structural questions and to propose a complete scenario of LFY evolution, we investigated the properties of LFY proteins from a hornwort and several algae species. We present the crystallographic structures of the promiscuous NaLFY from hornworts bound to type II and type III DNA sequences and the algal *Interfilum paradoxum* IpLFY bound to type III DNA. Type III binding reveals new protein DNA contacts and a novel DNA-binding mode: in this mode the two LFY-DBDs assemble on opposite sides of the DNA with dimer binding most likely due to DNA-mediated co-operativity and not direct protein-protein interactions as observed for type I and II sequences (this work and (Sayou et al. 2014)). Integrating these new findings with previous studies, we propose a more complete evolutionary scenario featuring two independent emergences of dimerization modes and multiple shifts in binding specificity.

## Results

### 1. Structural characterization of NaLFY-DBD bound to type II sequence

To investigate how the promiscuous NaLFY-DNA-binding domain (Na-LFYDBD) recognizes all three types of DNA motifs (Sayou et al. 2014), we determined its crystallographic structure in complex with type II DNA at 2.94 Å resolution (Fig. 1 and Table 1). The NaLFY-DBD/type II DNA complex adopts an architecture nearly identical to those of PpLFY-DBD/type II DNA (RMSD = 0.805 Å, PDB 4BHK) and AtLFY-DBD/type I DNA (RMSD = 1.652 Å, 2VY1 and 2VY2) (Sayou et al. 2014). All LFY DBD structures experimentally characterized to date adopt an all alpha-helical fold with both DBD monomers positioned on the same DNA face and engaging bases in the major and minor grooves (Fig. 1C-F). Previous studies also identified residue 312 as crucial for sequence-specific DNA interactions (Fig. 1D): this residue is an aspartate (D) in *Physcomitrium patens* LFY (PpLFY), a histidine (H) in *Arabidopsis thaliana* LFY (AtLFY), and a glutamine (Q) in NaLFY-DBD. The crystal structure reveals that glutamine 312 (Q312) in NaLFY-DBD forms a direct hydrogen bond with deoxycytosine (dC5) in the DNA major groove (Fig. 1E), analogous to the interaction between aspartate 312 (D312) and dC5 in PpLFY-DBD. This conserved interaction explains why NaLFY-DBD and PpLFY-DBD both bind the type II motif, despite their divergent evolutionary lineages.

**Table 1.** Data collection and refinement statistics.

|  | 8S8Q (IpLFY) | 9FWY(NaLFY II) | 9G19 (NaLFY III) |
| --- | --- | --- | --- |
| **Data collection** |  |  |  |
| Space group | P6<sub>1</sub> | P2<sub>1</sub>2<sub>1</sub>2<sub>1</sub> | P2<sub>1</sub>2<sub>1</sub>2 |
| Cell dimensions |  |  |  |
| *a*, *b*, *c* (Å) | 144.9, 144.9, 69.8 | 43.2, 90.0, 152.4 | 77.7, 154.1, 53.1 |
| $\alpha$, $\beta$, $\gamma$ (°) | 90, 90, 120 | 90, 90, 90 | 90., 90., 90. |
| Resolution (Å) | 78.1-2.94 (3.04-2.95)* | 78.-2.94 (3.012-2.94) | 77.-3.09 (3.2-3.09) |
| *R*<sub>sym</sub> or *R*<sub>merge</sub> (%) | 9.6 (160) | 15 (60.4) | 10.6 (104) |
| *I* / $\sigma I$ | 8.3 (2.0) | 2.38 (2.0) | 7.4 (2.0) |
| Completeness (%) | 94.9 (88.1) | 95 (98.3) | 98 (79.2) |
| Redundancy | 2.7 (2.8) | 2.7 (2.8) | 5.3 (5.3) |
| CC(1/2) | 99.6 (47.1) | 99.5 (82.9) | 99.8 (80.9) |
| **Refinement** |  |  |  |
| Resolution (Å) | 47.-2.95 (3.0-2.95) | 20.-2.94 (3.01-2.94) | 30.-3.09 (3.17-3.09) |
| No. reflections | 16920 (1246) | 12200 (904) | 10489 (607) |
| *R*<sub>work</sub> / *R*<sub>free</sub> | 19.8/23.8 (47./49.2) | 25.7/29.7 (38./41.) | 24.3/30.3 (34.5/35) |
| No. atoms | 3602 | 3777 | 3456 |
| Protein | 2599 | 2575 | 2438 |
| DNA | 985 | 1185 | 1017 |
| Water | 30 | 17 | 1 |
| Other ligands | - | - | - |
| *B*-factors |  |  |  |
| Protein | 55. | 33.6 | 215. |
| DNA | 65. | 44.3 | 206. |
| Water | 79. | 59.5 | 300. |
| Other ligands | - | - | - |
| R.m.s. deviations |  |  |  |
| Bond lengths (Å) | 0.005 | 0.01 | 0.01 |
| Bond angles (°) | 1.7 | 1.36 | 1.04 |

\* Refers to the highest resolution shell

This structural conservation also underscores a shared dimerization interface, even as DNA-binding specificity diverged. While the DBDs of NaLFY, AtLFY, and PpLFY all homodimerize through the H387 residue on helix α7, NaLFY features a slightly distinct dimerization interface. Specifically, the R390 residue—present in AtLFY and PpLFY and known to contribute to dimerization—is replaced by K390 in NaLFY, which does not participate in a direct protein-protein interaction based on the crystal structure. Instead, the NaLFY DBD dimerization interface includes a different interaction: the glutamic acid, E364, forms a hydrogen bond with the backbone amide nitrogen of cysteine, C286, on the partner monomer (Fig. 1D). As we were unable to obtain a structure of NaLFY-DBD bound to type I DNA, we examined an AlphaFold3 model that retains dimer and DNA-binding interactions with a potential additional interaction between Q312 and the phosphate backbone, suggesting an energetically favorable binding mode for the type I DNA motif (Supp. Fig. 1A).

### 2. Structural characterization of NaLFY bound to type III sequence

Next, we obtained the structure of the NaLFY-DBD bound to the type III DNA motif at 3.09 Å resolution (Fig. 2A, Table 1). Each NaLFY monomer exhibited the same fold as previously described above, but with a new orientation of two LFY monomers bound to the type III DNA sequence. The contacts between the protein and DNA closely resemble those observed in the type II DNA complex (Fig. 2C-D). However, the relative positioning of the NaLFY-DBD monomers on the DNA markedly diverges from the previously characterized type I and type II LFY structures (Fig. 2A-B). Here, the two NaLFY-DBD monomers bind to opposite sides of the DNA, with no direct protein-protein contacts (Fig. 2A).

**Figure 2:**
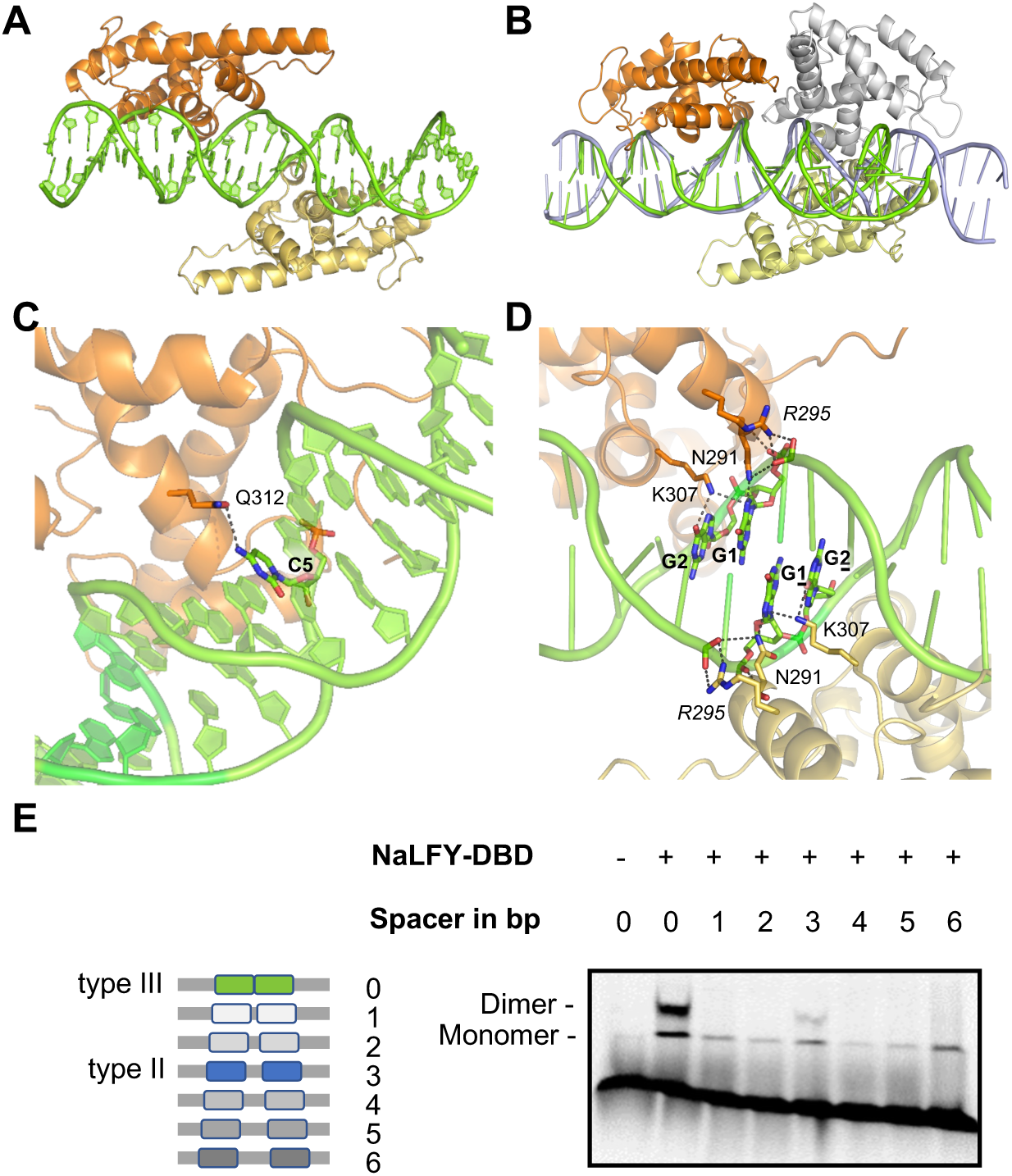
Structural basis of promiscuous NaLFY-DBD for type III specificity. (A) Crystal structure of NaLFY-DBD (yellow and orange) bound to type III DNA (green). (B) Comparison between NaLFY binding to type II (blue) and III (green) DNAs. One monomer used for alignment is shown as an orange cartoon, the two other ones are shown in yellow (type III) and grey (type II). Note the very different DNA shapes associated to each configuration. (C) Close-up on specificity determinant residues (Asparagine, N292) and DNA bases (desoxycytosine, C; desoxyguanine, G). (D) Close-up view showing how each monomer contacts G1G2 bases on one half site and the phosphate backbone of C1 from the other half site. Interacting residues are N291, R295 and K307, black residues interact with bases and red with phosphate backbone. (E) EMSA assaying NaLFY-DBD (5 µL of soluble fraction) binding to various half sites (represented as a colored box) separated by 0 to 6 bases.

The absence of a protein-protein interaction surface to mediate dimerization raises the question: Why do two NaLFY-DBDs preferentially bind half-sites spaced by 0 bp? To address this, we hypothesized that DNA-mediated cooperativity—rather than protein-protein interactions—stabilizes the dimeric complex. Using Electrophoretic Mobility Shift Assays (EMSA) with DNA probes containing variable spacer lengths between LFY half sites, we found that NaLFY-DBD indeed exhibits a marked preference for half sites separated by 0 bp (as in type III sites) or 3 bp (as in type II sites, weaker than for type III), but not for other spacer configurations (Fig. 2E). Using increasing protein concentrations, we further demonstrated the highly cooperative binding of NaLFY-DBD to DNA with half sites separated by 0 bp as compared to 6 bp spacers (a distance chosen to ensure the absence of contacts between monomers) (Supp. Fig. 1B). These data suggest that dimer assembly is indeed stabilized by DNA-mediated cooperativity (Morgunova and Taipale 2017), wherein the binding of one monomer deforms the DNA, facilitates recruitment of the second monomer. This mechanism is supported by the pronounced DNA bending observed in the type III complex when compared to the type I and II bound complexes (Fig. 2B). It suggests that binding of a first DBD monomer may indeed induce structural alterations at the central C1G1 bases (5’-TTGCTAC**<u>CG</u>**GTCGCTG-3’, Fig. 2D) that are contacted by residues N291 and R295 from each DBDs. These alterations likely lower the energetic barrier for the recruitment of the second monomer, thereby promoting cooperative binding.

Thus, NaLFY demonstrates remarkable versatility in dimerization, adopting two distinct dimeric conformations on DNA: one stabilized by direct protein-protein dimerization (as observed previously for PpLFY and AtLFY) and another facilitated by DNA-mediated co-operativity.

### 3. Characterization of algal LFY DBDs bound to DNA

To further explore the DNA-binding properties of LFY proteins, we also investigated a diverse set of charophyte algae. Previous work had characterized KsLFY, the LFY protein from *Klebsormidium subtile*, revealing its strict dimeric binding to the type III DNA motif (Sayou et al. 2014). However, it remained unknown whether this behavior is conserved across other algal LFY proteins. To address this question, we systematically mined available transcriptomic and genomic resources within this plant group, focusing on *LFY* sequences containing a complete DBD. This effort led to the identification of numerous new *LFY* genes spanning all charophyte clades. We also identified a highly divergent and likely partial LFY sequence from the early-diverging alga *Spirotaenia* (Supp. Fig. 2)—for which we found no evidence of DNA binding. Interestingly, the discovery of this gene extends the origin of the *LFY* gene to the very base of streptophyte plants, predating earlier estimates (Gao et al. 2019). However, the properties of this protein remain unknown.

We selected representative LFY proteins from each major class—CcLFY from *Cylindrocystis cushleckae*, CiLFY from *Coleochaete irregularis*, CvLFY from *Chara vulgaris*, and IpLFY from *Interfilum paradoxum*—and assessed the DNA-binding preferences of their DBD across motifs with variable spacer lengths using band shift assays (Fig. 3). Strikingly, the strongest binding was consistently observed with the 0-bp spacer configuration, supporting that the DNA-mediated cooperativity observed for NaLFY-DBD applies to all these proteins. Unlike KsLFY-DBD, however, these LFY proteins also demonstrated the ability to bind DNA as monomers, albeit with a reduced affinity (in the case of CvLFY-DBD) and even as a dimer with an alternate half-site spacing (as CiLFY-DBD with a 1-bp spacer). We refer to these binding preferences (monomeric, and dimeric on the 0-bp spacer) as ‘relaxed type III’ specificity. This binding mode differs from the promiscuous feature of NaLFY-DBD as the algal proteins IpLFY and CiLFY cannot bind type I and II sites as dimers (Supp. Fig. 3). Notably, the dimerization on the 3-bp spacer DNAs occurs with NaLFY-DBD but not with these algal LFY-DBDs, consistent with the absence of the residues necessary for protein-mediated dimerization.

**Figure 3:**
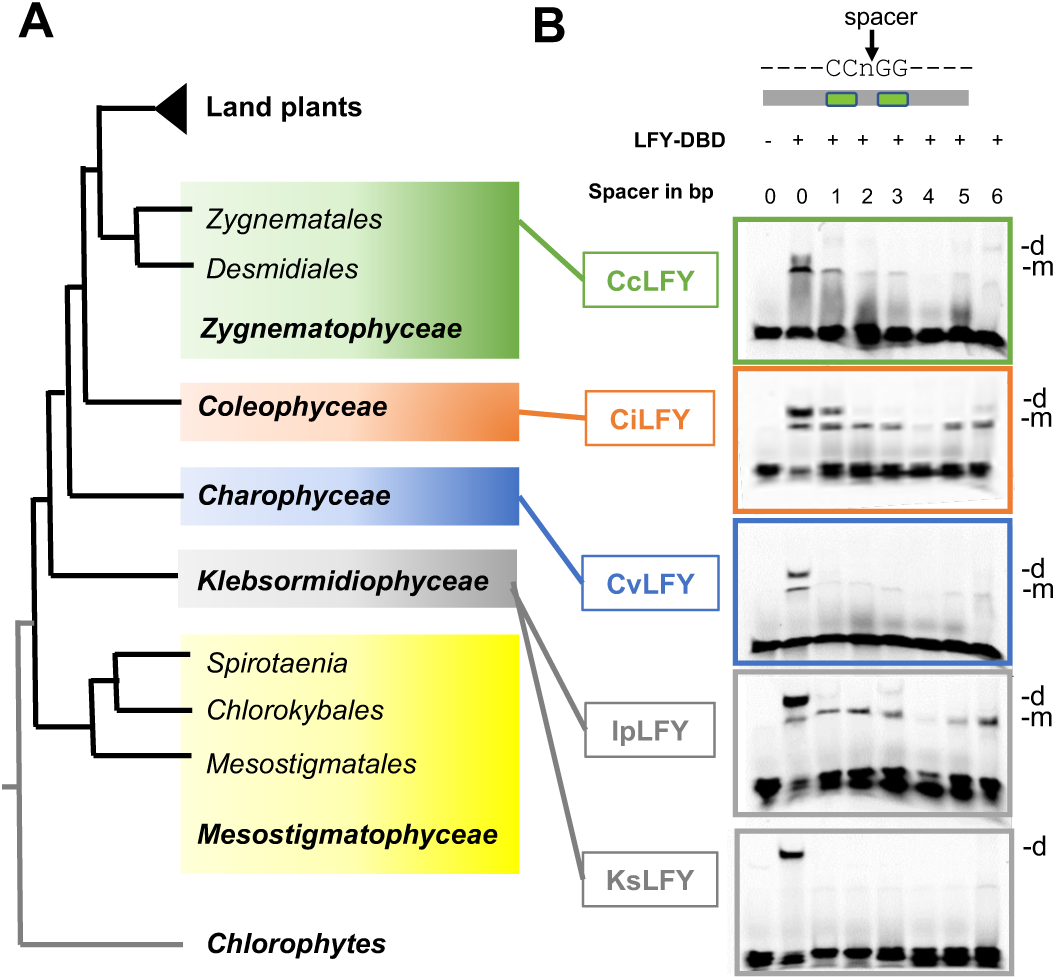
Charophyte LFY-DBDs prefer a 0-bp spacer motif. A) Simplified phylogenetic species tree showing the groups of charophyte algae. B) EMSA testing the ability of LFY-DBD from various charophyte species to bind half sites (taken from a type III site) spaced by a variable number of base pairs (0 to 6): *Cylindrocystis cushleckae* (CcLFY). *Coleochaete irregularis* (CiLFY), *Chara vulgaris* (CvLFY), *Interfilum paradoxum* (IpLFY) and *Klebsormidium subtile* (KsLFY).

To further elucidate the structural properties of LFY from early-diverging charophytes, we solved the crystallographic structure of IpLFY-DBD in complex with type III DNA at 2.95 Å resolution (Fig. 4, Table). The overall binding configuration closely resembles that of NaLFY-DBD on the same type III DNA (Fig. 4, Fig. 2B). The N291 residues from each monomer interact symmetrically to sandwich the DNA, as previously described for NaLFY-DBD (Fig. 4B). The DNA conformation induced by IpLFY is also highly similar to that observed with NaLFY, suggesting that DNA-mediated cooperativity may operate through comparable mechanisms in both proteins studied to favor binding to two binding sites spaced by 0 bp (Supp. Fig. 3).

**Fig. 4:**
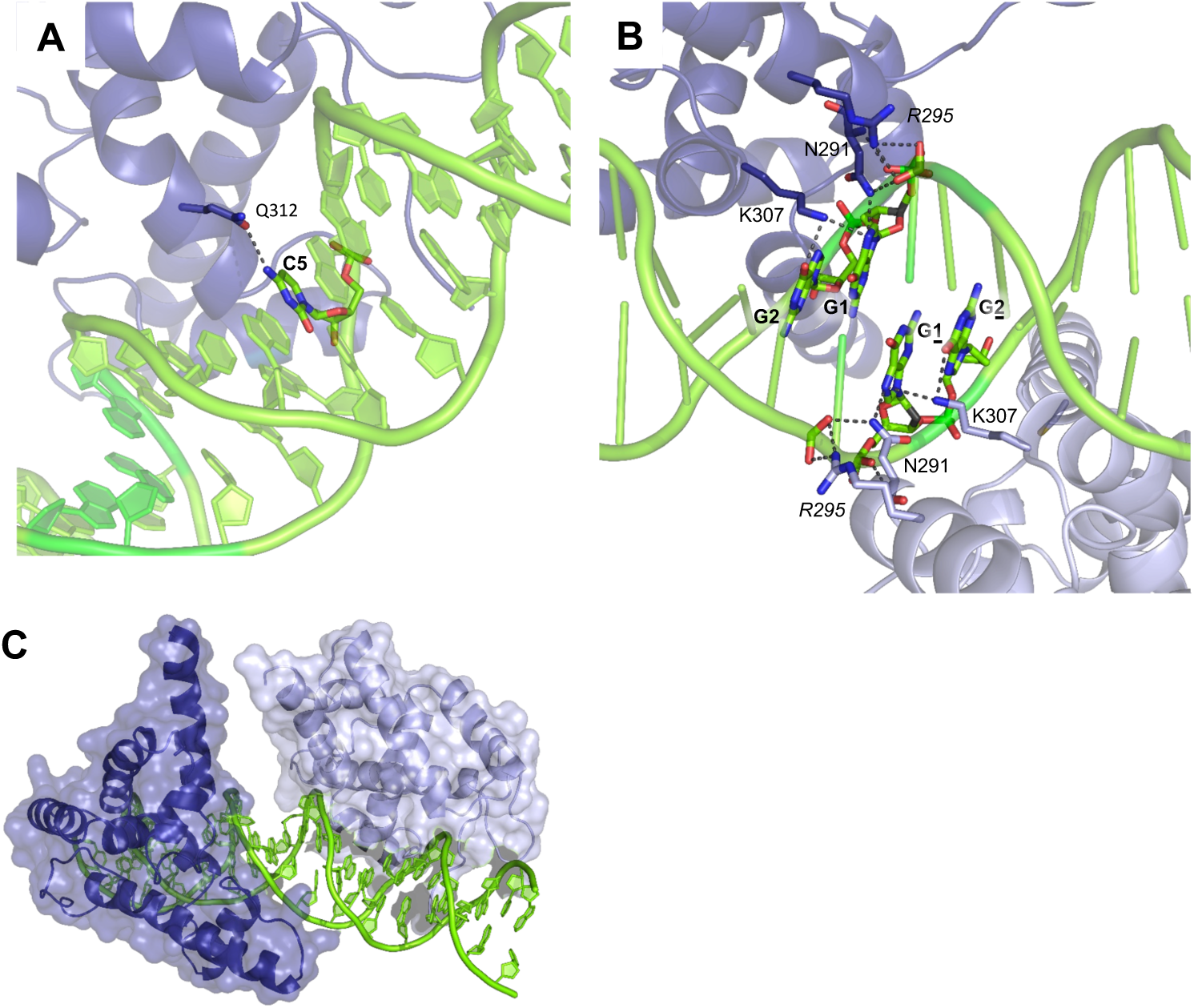
Structural basis of IpLFY-DBD for type III specificity. Crystal structure of IpLFY-DBD (blue) bound to type III DNA (green). (A) Close-up on interaction of Q292 and DNA bases (desoxycytosine, C; desoxyguanine, dG). (B) Close-up on residue involved in DNA contact in IpLFY-DBD (C) Surface representation of IpLFY on type III DNA illustrating the absence of contacts between the two monomers.

In an effort to understand the strictly dimeric behavior of KsLFY and after failing to obtain its crystallographic structure, we employed AlphaFold3 (AF3) to model the KsLFY/DNA complex (Abramson et al. 2024). The predicted KsLFY/DNA structure was nearly identical to the experimentally determined IpLFY/DNA structure, offering no clear explanation for the obligate dimeric binding observed with KsLFY (Supp. Fig. 4A). We suspected that the extended C-terminal tail of KsLFY might contribute to its dimerization properties, but neither our experimental data nor the AF3 model provided compelling support for this hypothesis (Supp. Fig. 4B). We thus could not conclude if the strict dimeric behavior of KsLFY DBD results from an unidentified protein interface of the C-terminal extension or unstable monomer on DNA.

### 4. Discussion

Since the initial evolutionary scenario proposed to account for LEAFY-DBD’s structural diversification (Sayou et al. 2014), significant advances in plant phylogeny—in particular regarding bryophytes and algae (Rensing 2018; Leebens-Mack et al. 2019; Donoghue et al. 2021; McCourt et al. 2023)—and the discovery of *LFY* genes across diverse species (Brunkard et al. 2015; Gao et al. 2019) have provided new opportunities to revisit this framework. Our goal here was to take advantage of these findings to elucidate the early evolution of LEAFY during the critical water-to-land transition and to construct an evolutionary narrative consistent with protein sequences, biochemical assays and additional experimentally determined crystal structures.

Here, we focus on the isolated LFY DNA-binding domain (DBD) to dissect its intrinsic DNA-binding properties and their evolution across plant species. It should be noted, however, within the full-length protein, the DBD is invariably associated with the Sterile Alpha Motif (SAM) oligomerization domain, which promotes higher-order complex formation and enhances DNA binding (Sayou et al. 2016). Depending on the complex examined, the presence of the SAM domain can have contrasting effects: it prevents the binding as monomer of AtLFY and *Ginkgo biloba* LFY to type I half sites (Sayou et al. 2016) but does not affect the capacity of an NaLFY monomer to bind on a type III half site (Sayou et al. 2014). Consequently, the properties of LFY full-length protein are potentially determined not only by the DBD itself but also by other domains. Moreover, in angiosperms, LFY (as monomer and homodimer) was shown to also bind DNA in complex with the F-box protein, Unusual Floral Organs (UFO) (Rieu et al. 2022). It remains unclear whether similar interactions occur in non-flowering plants. In this study, we have specifically examined the DNA-binding properties of the isolated LFY DBD, independent of its broader protein context or potential interacting partners. This targeted approach allows us to dissect the intrinsic DNA-binding capabilities of the DBD, providing a foundation for understanding its evolutionary trajectory (structures available are gathered on Supp. Fig. 5).

#### A structural scenario for LFY evolution

The results presented here provide new insights into the early stages of LEAFY evolution. First, we identified a novel and highly divergent LFY form in the streptophyte alga Spirotaenia, suggesting that LFY originated at the very base of the streptophyte lineage—earlier than previously proposed (Gao et al. 2019) (Fig. 5).

**Figure 5:**
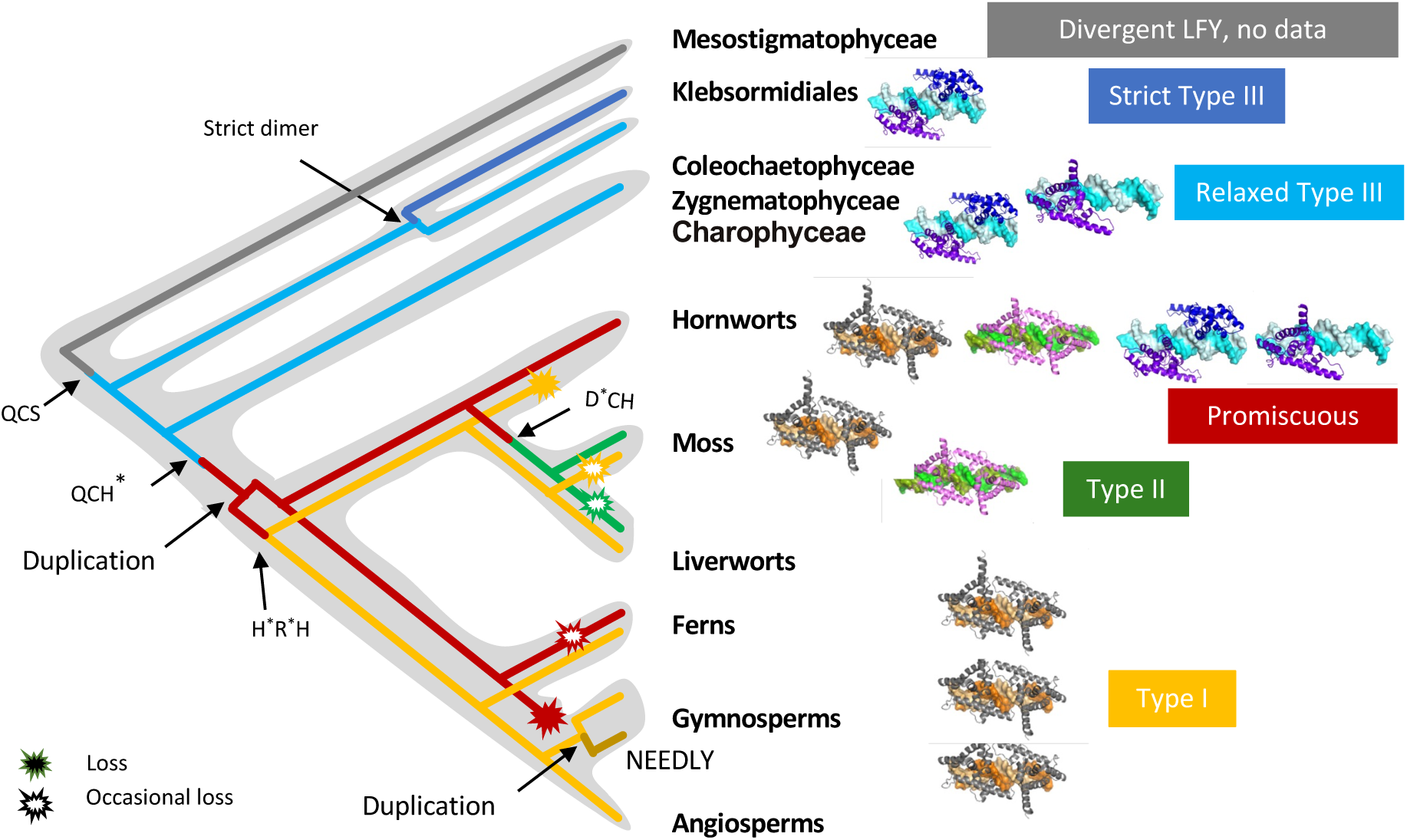
Scenario for LFY-DBD evolution. Gene duplication events are highlighted within the grey-shaded regions. Asterisks (*) denote the changes affecting functional residues, marking four key evolutionary events. Complete gene loss is indicated by solid symbols, while occasional loss is represented by open symbols.

Second, our analysis of LFY-DBDs from various charophyte algae support the hypothesis that early LFY-DBD prefers to bind to DNA as dimers, through DNA-mediated cooperativity. Indeed, in the LFY- type-III-DNA complexes, the absence of direct contact between the two LFY-DBD, combined with the observation that both monomers interact with shared bases within a deformed DNA environment, strongly suggests that the dimeric preference arises from DNA-mediated cooperativity. However, whereas we originally proposed a strict dimeric binding to type III DNA based on KsLFY-DBD, it appears that charophytes LFY indeed exhibit a preferential dimeric binding of type III DNA but are also capable of binding as monomers—"relaxed type III" specificity. Based on these findings, the most parsimonious hypothesis is that the relaxed type III specificity represents the ancestral state, while the strict dimeric type III specificity observed in KsLFY is likely derived. However, we cannot definitively exclude the possibility that the strictly dimeric binding of KsLFY reflects the ancestral condition.

Integrating our structural findings with current phylogenetic models, specifically the bryophyte monophyly hypothesis (Donoghue et al. 2021; Chen and de Vries), we propose that LFY emerged at the base of streptophytes. This scenario posits that early LFY-DBDs exhibited relaxed type III specificity, binding DNA cooperatively via DNA-mediated interactions. The subsequent evolution of protein-protein dimerization (e.g., via the H387 residue) likely enabled the transition to type I/II motifs, facilitating the diversification of LFY’s regulatory roles during land colonization. In charophytes, LFY-DBDs display a relaxed type III specificity and a conserved glutamine at position 312 in direct contact with the C5 base. The residues that, in land plants, mediate protein-protein dimerization are absent (Fig. 5). A key evolutionary shift likely occurred during the water-to-land transition with the appearance of a histidine at position 387 in helix 7. This substitution enabled DNA-binding domain dimerization via direct protein-protein interactions, allowing a novel dimerization mode. As H387 is present in all LFY copies in land plants, the novel dimerization mode likely preceded the proposed LFY gene duplication, also thought to have occurred around the origin of land plants (Gao et al. 2019). The new DNA-binding conformation, featuring two interacting DBDs positioned on the same side of the DNA, was fixed and retained during the subsequent shift in specificity toward type I or type II motifs. Such a configuration might present some specific selective advantage. Indeed, studies in Arabidopsis have proposed that LFY could work as a pioneer transcription factor, able to bind to and open closed chromatin regions (Sayou et al. 2016; Jin et al. 2021; Lai et al. 2021). Binding to nucleosomal DNA is likely facilitated if the two LFY monomers sites are located on the same (accessible) side of the DNA. We therefore speculate that this novel configuration may have enabled LEAFY to acquire its proposed pioneer transcription factor properties, although this remains to be further investigated.

After LFY gene duplication, one the two duplicates persisted as promiscuous in hornworts and some ferns but was lost in other ferns and seed plants (Fig. 5). In mosses, this duplicate underwent a glutamine (Q) to aspartate (D) substitution, leading to the type II specificity unique to mosses. The second duplicate experienced a glutamine-cysteine (QC) to arginine-histidine (RH) transition, resulting in the type I specificity that persisted in liverworts, some mosses, and all vascular plants, but was lost in hornworts and some moss species. Overall, this scenario illustrates a shift in dimerization mode from DNA-mediated cooperativity to direct protein-protein interaction, and a transition through a promiscuous intermediate that likely facilitated the gradual changes in binding specificity, as previously proposed. The duplication of the promiscuous form, now well supported by the identification of numerous LFY sequences (Brunkard et al. 2015; Gao et al. 2019), provided additional flexibility for this low-copy and conserved transcription factor to evolve new specificities over time.

Under the bryophyte monophyly hypothesis, this scenario involves four changes in key residues (QCS → QCH*, QCH → H*R*H and QCH → D*CH, Fig. 5) and two duplications. It can be adapted to alternative phylogenetic models, some of which necessitate even fewer changes. However, the core principles of our model—combined promiscuity as a transitional state and gene duplication facilitating functional diversification—remain robust across different phylogenetic contexts.

Despite this progress, several key questions remain unresolved. The DNA-binding specificities of SpiroLFY and NDLY from gymnosperms (Moyroud et al. 2017) have yet to be characterized, leaving gaps in our understanding of LFY’s functional diversity across early-diverging lineages. Additionally, the rarity with which LFY gene duplicates are maintained throughout evolution is striking, with diversification events occurring only in isolated cases, such as in mosses and gymnosperms (Gao et al. 2019). The reasons for the apparent detrimental effects of maintaining additional LFY copies remain unclear and warrant further investigation. Finally, the evolutionary origin of this unique DNA-binding domain, which bears little resemblance to other known protein folds, remains an intriguing and open question.

In conclusion, this study not only elucidates the evolutionary trajectory of the LEAFY transcription factor but also serves as a broader model for understanding the structural and functional diversification of transcription factors. By revealing how protein-DNA interactions evolve and adapt across different lineages, our findings offer valuable insights into the mechanisms driving the evolution of gene regulatory networks. Ultimately, this work underscores the dynamic interplay between protein structure, DNA-binding specificity, and gene duplication in shaping the functional diversity of transcription factors over evolutionary time.

## Methods

### *LFY* genes identification

To better understand the origin and evolution of LFY, we searched for *LFY* sequences in several databases, including NCBI (version 5) and the OneKP Project database (version 5, (Matasci et al. 2014); 1KP initiative(Leebens-Mack et al. 2019)). As a query, we used the LFY-DBD protein sequences previously identified in early-diverging green algae, such as KsLFY or CvLFY(Sayou et al. 2014). We used standard parameters with filter restricting the finding in green algae lineage data. Using this approach, we identified several previously reported LFY protein sequences, including the LFY sequences described in the genus *Interfilum* (e.g. scaffold ID: FPCO_scaffold_2028225), *Mesotaenium* (e.g. scaffold ID: NBYP_scaffold_2007098) or *Chaetosphaeridium* (e.g. scaffold ID: DRGY_scaffold_2003051)(Gao et al. 2019). In the *Spirotaenia* transcriptome, we also detected putative LFY-like sequences (SpiroLFY) distributed across three different scaffolds (IDs: TPHT_scaffold_2002904, NBYP_scaffold_2007097, TPHT_scaffold_2002906, and TPHT_scaffold_2002903). These sequences include a primitive LFY-DBD DNA signature representing approximately 40% sequence similarity to typical algal LFY-DBDs protein (on 137 putative amino acid sequence blasted using IpLFY-DBD as a query for example). Interestingly, the α3 helix of the LFY-DBD, which is crucial for DNA-binding activity (Hamès et al. 2008), is partially conserved in these SpiroLFY putative sequences (with a perfect match for the “CYALHCLD” amino acid putative sequence of the helix).

### Cloning and production of LEAFY DBDs

Partial cDNA sequences from algae LFY corresponding to the DNA binding domains were cloned using a Gibson assembly strategy in the pETM-11 vector (Dümmler et al. 2005) to add a 6xHis tag. Briefly, pETM-11 vector was digested using the NcoI-XhoI restriction sites and cDNA amplicon corresponding to the LFY DBDs were introduced to obtain the following expression clones: pHC191 (*KsLFY-DBD*), pHC192 (*IpLFY-DBD*), pHC193 (*NaLFY-DBD*), pHC194 (*CvLFY-DBD*), pHC195 (*CcLFY-DBD*), pHC196 (*CiLFY-DBD*). Oligonucleotides and details on the cloning strategy are provided in Table S1

The pETM-11-based expression vectors were introduced in *Escherichia coli BL21* cells strain (Novagen). Cell cultures were grown at 37 °C to reach OD_600_ of 0.6, then the cultures were cooled at 18 °C and protein expression was induced by adding 1 mM of IPTG for 12 h. Cultured cells were collected by centrifugation and the pellet was resuspended in Buffer A (Tris-HCL 20 mM pH 8, TCEP 1 mM) supplied with complete protease inhibitors (Thermo). Cells were lysed by sonication, and the soluble fraction was loaded onto a denaturing polyacrylamide gel.

### Electrophoretic mobility shift assay (EMSA)

All oligonucleotides used are listed in Supplementary Table S1. EMSAs were performed using 10 nM DNA labelled with Cy5 as described (Moyroud et al. 2011). Recombinant proteins were incubated with labelled DNA in the Binding buffer (BB) (HEPES 10 mM pH 7.5, Spermidine 1 mM, EDTA 14 mM, BSA 0.3 mg/mL, CHAPS 0.25 %, glycerol 2 %, fish sperm DNA 28 ng/mL, TCEP 3 mM) with 250 mM NaCl final concentration, and incubated for 15 min at 4 °C in the dark. The reaction was loaded on a native 5 % acrylamide gel, electrophoresed at 90 V for 1.5 h at 4 °C, and scanned on ImageQuant 800 (Cytiva).

Recombinant proteins were produced in *Escherichia coli* Rosetta TM 2 (DE3) strain (Novagen). A pellet corresponding to 20 mL of culture was resuspended in 3 mL of buffer containing Tris-HCl 20 mM, DTT 1 mM, protease inhibitor (Pierce). After sonication, a centrifugation is realized to separate soluble fraction. EMSA were done using 5 µL of this soluble fraction. Reactions were performed in the BB buffer as previously described.

### Crystallization of LFY-DNA complexes

HPLC purified ssDNA oligonucleotides were ordered from Eurofins and used without further purification. Equimolar concentrations of the two oligomers were mixed in LFY purification buffer, heated to 100°C for 5 min and annealed on the benchtop overnight. Purified LFY protein was mixed with a 1:1.1 ratio of annealed dsDNA to form the LFY-DNA complex. Crystallization experiments were carried out by the vapour-diffusion method at 4°C using sitting drops with a 1:1 ratio of protein–DNA complex:precipitant with a final protein concentration of ∼5-10 mg.ml^−1^. Crystallization was performed by the EMBL High Throughput Crystallization Facility, Grenoble, France. All crystallization experiments were performed in 200 nl sitting drops using a Cartesian PixSys 4200 crystallization robot (Genomic Solutions, UK) using Greiner CrystalQuick plates (flat bottom, untreated) and imaged with a Rock Imager (Formulatrix, USA). Suitable well diffracting crystals were grown after ∼ 1 week in 50 mM sodium acetate, pH 4.6, 8 % PEG 3350 and cryoprotected with 20 % glycerol (IpLFY), 50 mM sodium acetate, pH 4.6, 20 % PEG3350, 100 mM ammonium chloride and cryoprotected with 20% glycerol (NaLFY+type II) or 50 mM MES, pH 6.8, 20% PEG3350 and 200 mM lithium chloride and cryoprotected with 20% glycerol (NaLFY+type III 9G19).

### Structure resolution, model building and refinements

Diffraction data were collected at 100 K at the European Synchrotron Radiation Facility, Grenoble, France, on ID23-2 at a wavelength of 0.873 Å. Indexing was performed using MXCube (Gabadinho et al. 2010) and the default optimized oscillation range and collection parameters used for data collection. All datasets were integrated and scaled using the programs XDS and XSCALE (Kabsch 2010). Molecular replacement for all datasets was performed with Phaser (McCoy 2007). Model building was performed using Coot (Emsley et al. 2010) and all refinements were carried out in Refmac (Murshudov et al. 1997). The structure quality was assessed using MolProbity (Williams et al. 2018) Data collection and refinement statistics are given in Table 1. The structure is deposited under PDB codes 9FWY (NaLFY type II), 9G19 (NaLFY, type III) and 8S8Q (IpLFY).

## Data availability

Structural data are deposited in the Protein Data Bank under accession codes 8S8Q, 9FWY, and 9G19. Oligonucleotide sequences are listed in Table S1. Raw data (e.g., EMSA gels) are available from the corresponding author upon request.

## Author contributions

FP designed the study, LV, ET, HC and LG performed biochemical experiments, LV and CZ performed structural experiments, CZ and MHN solved the structures, HC performed LFY searches, FP, RD, CZ analyzed the results and supervised the work, FP and CZ wrote the manuscript with the help of all authors.

## Acknowledgments

We gratefully acknowledge the European Synchrotron Radiation Facility (ESRF) for granting beamtime on the ID23 beamline.

## Declaration of generative AI and AI-assisted technologies in the manuscript preparation process

During the preparation of this work the authors used Le Chat by MISTRAL AI in order to polish the English.

## Supplementary Information

**Figure S1:**
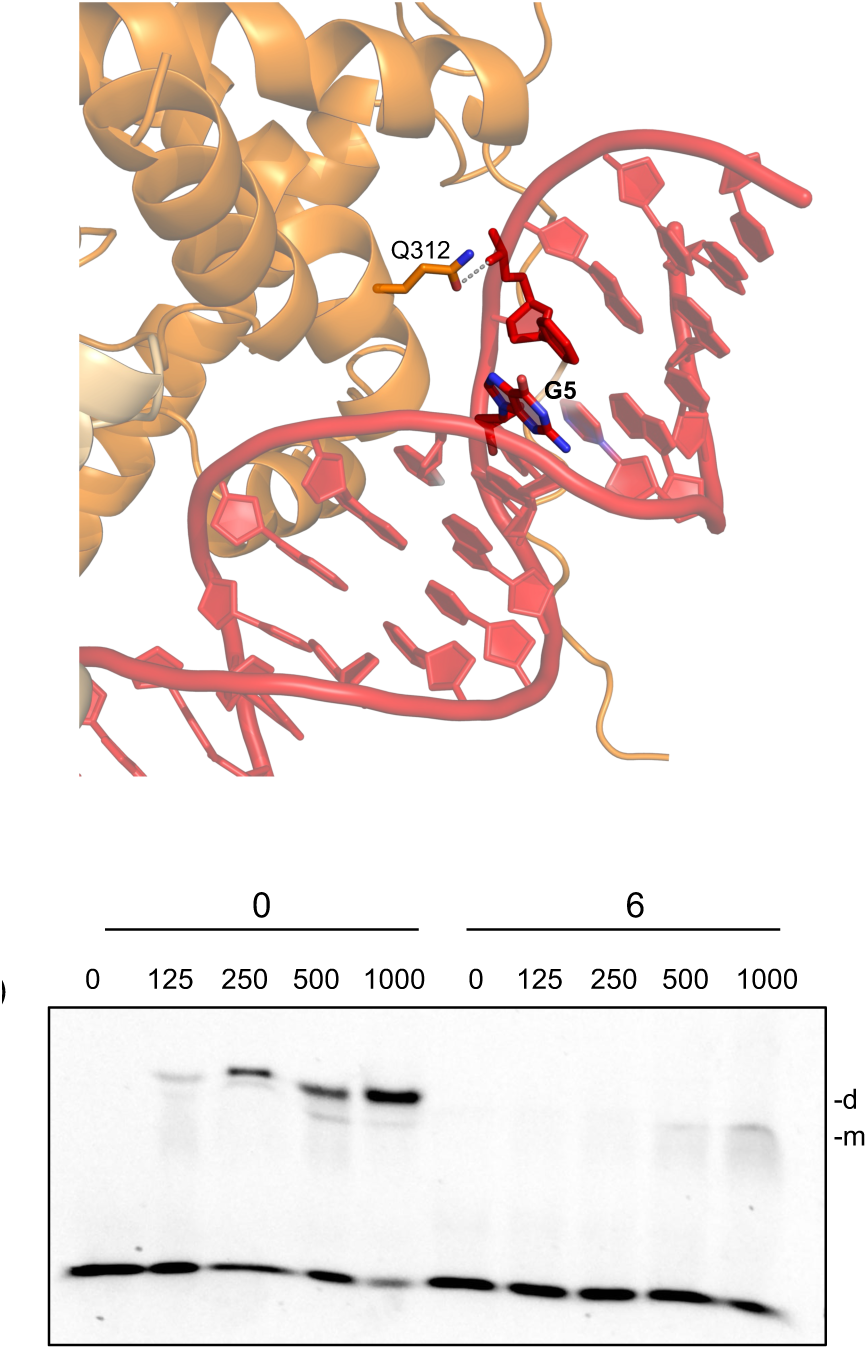
NaLFY-DBD characterization. A) Alphafold3 model of NaLFY-DBD on type 1 DNA model (sequence) showing that Q312 interacts with the phosphate backbone. B) EMSA showing the cooperative binding of NaLFY-DBD to DNA with 0-bp spacer as compared to 6-bp. On the 0-bp spacer, the dimeric complex is always more intense than the monomeric, even at the lowest concentrations. In contrast, binding to the 6-bp spacer is weaker and mostly monomeric.

**Figure S2:**
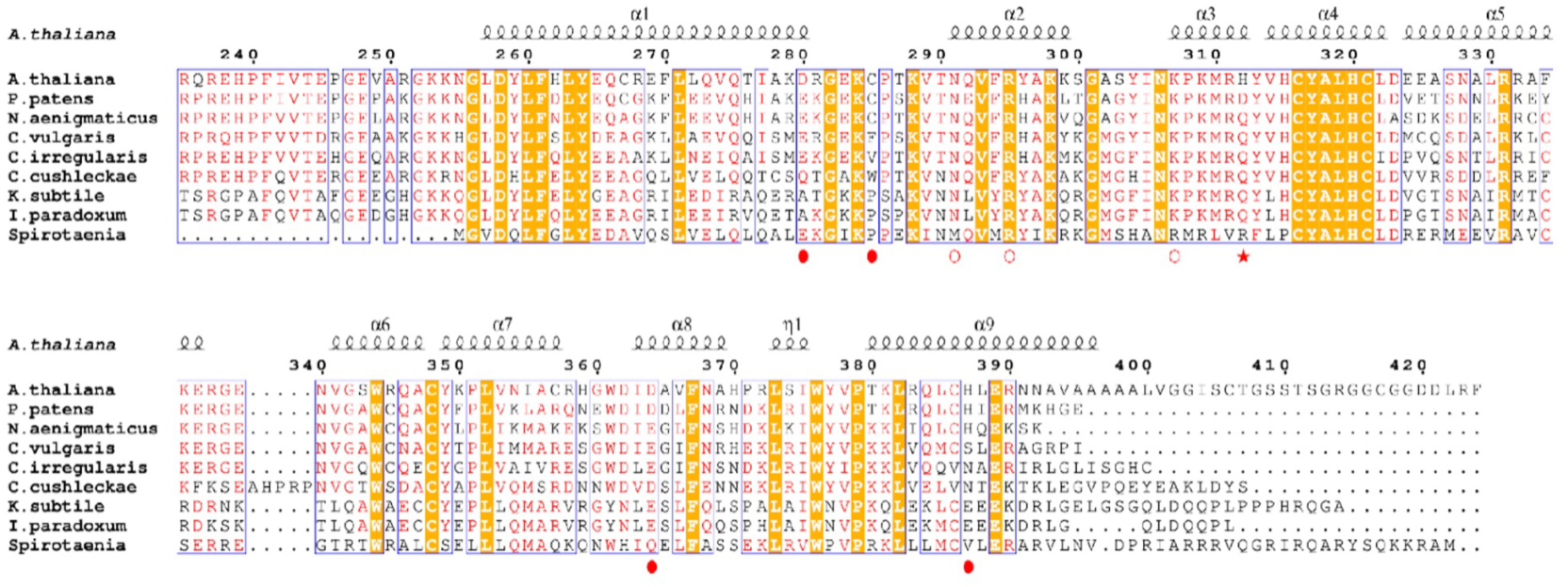
LFY-DBD sequence alignment. Alignment of LFY-DBD protein sequences including the divergent Spirotenia LFY. The plain circles indicate residues involved in DBD homodimerization in type I and II motifs, the red star indicates the residue interacting with DNA bases in type II and III motifs, open circle indicate residues interacting with the central CG bases in motif III.

**Figure S3:**
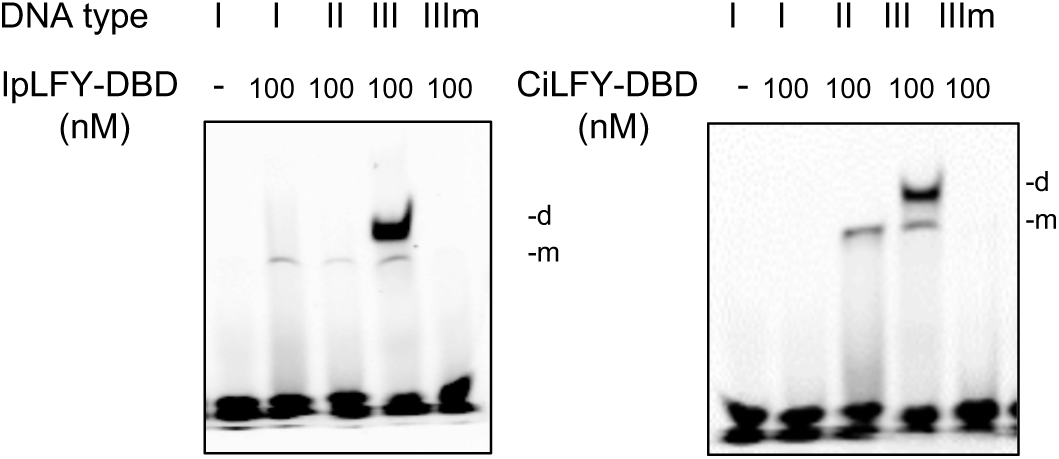
IpLFY-DBD and CiLFY-DBD algal proteins are not promiscuous. EMSA showing that IpLFY-DBD and CiLFY-DBD do not bind type I and type II DNA as dimers as NaLFY does. Type IIIm DNA has 2 mutations as compared to type III DNA (A_3_C_2_C_1_ replaced by C_3_C_2_A_1_). 100 nM of recombinant DBDs were used.

**Figure S4:**
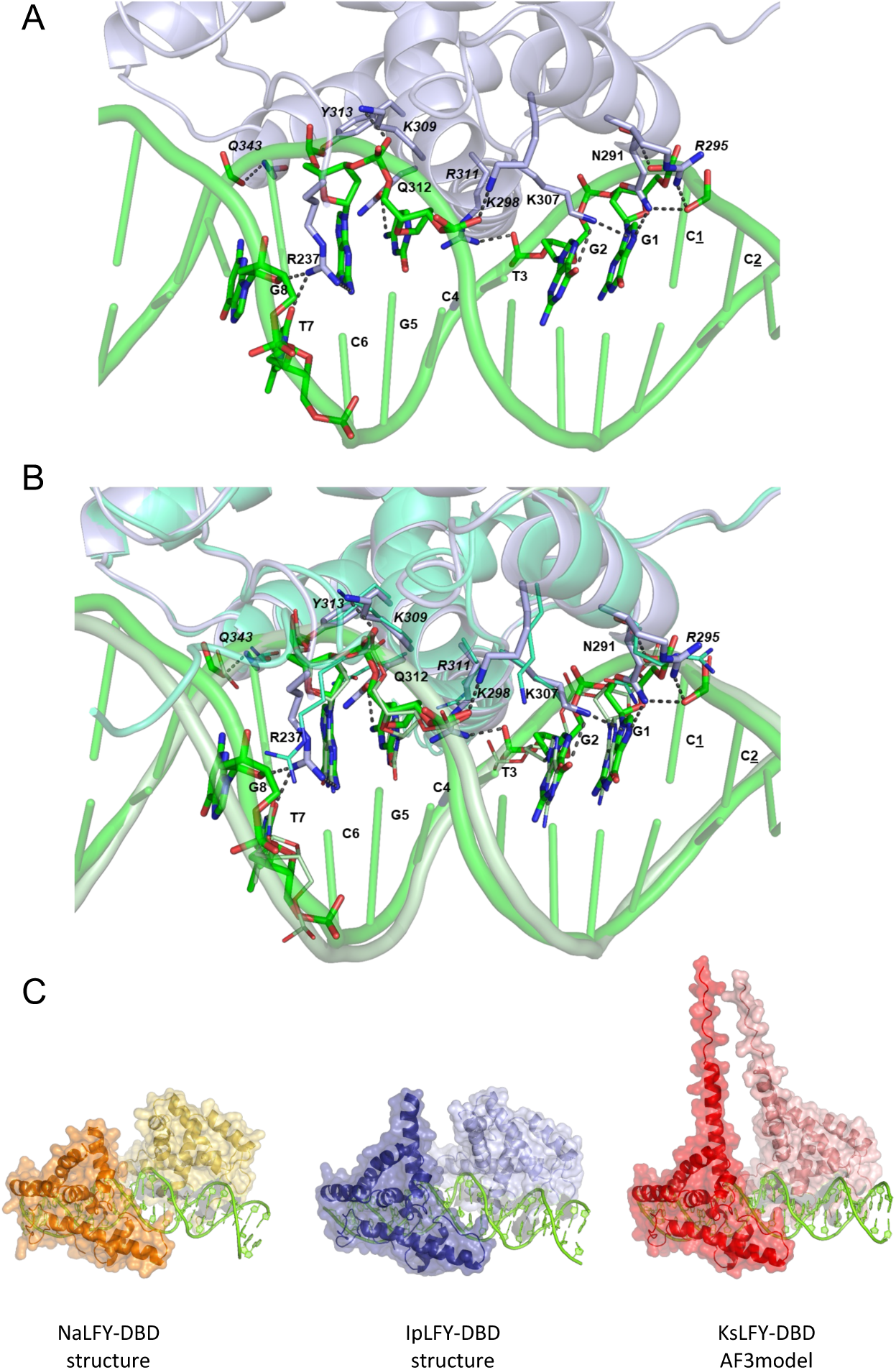
Alphafold3 KsLFY-DBD model comparisons. A. Structure of IpLFY-DBD monomer on type III DNA. B. Superimposition of Alphafold3 KsLFY-DBD model (light blue, thin sticks) and IpLFY-DBD structure (grey) in complex with type III DNA showing that both proteins exhibit similar interaction with DNA. Italic residues interact with the phosphate backbone. Other residues interact with DNA bases. Nucleotides are indicated on one strand. Only interactions below 3.4 Å are shown. C. Comparison of NaLFY-DBD, IpLFY-DBD structures and Alphafold3 KsLFY-DBD model in complex with type III DNA. Despite displaying a longer C-terminal sequence, KsLFY-DBD is not predicted to form protein dimerization interface.

**Figure S5:**
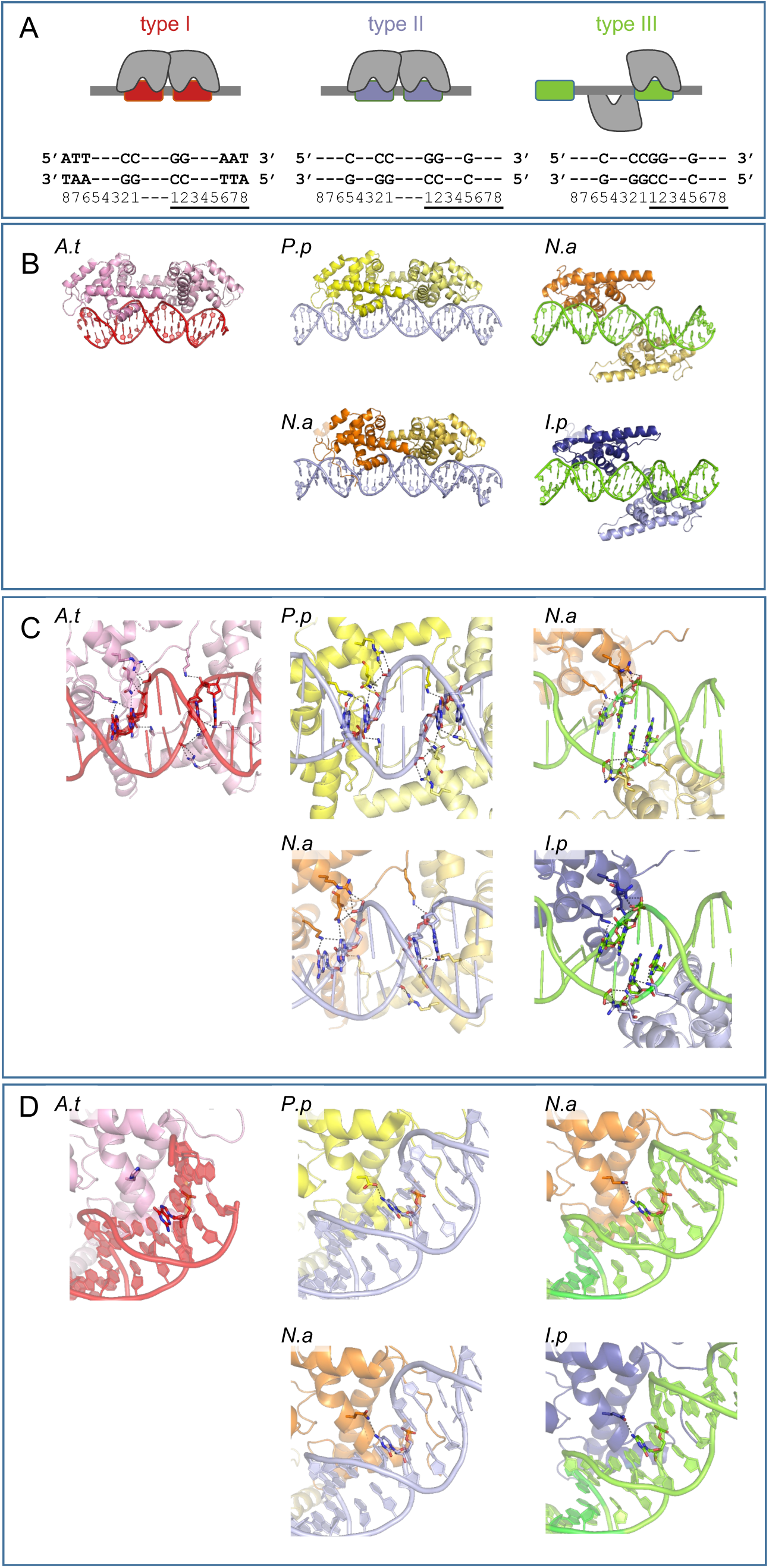
Summary of structures. A) Schematic representation of binding site providing the DNA color code. B) Overall position of the two monomers on DNA in the various configurations. C) View showing the interaction with the C1C2 from the major grove. D) View showing the interaction with C3. Data were taken from this work and Species are indicated by first letters *A.t* = *Arabidopsis thaliana*, *P.p* = *Physcomitrium patens*, *N.a*. = *Nothoceros aenigmaticus*, *I.p* = *Interfilum paradoxum*

## Supplementary methods

Pellet containing LFY protein was resuspended in Buffer A (Tris-HCl 20 mM pH8, TCEP 1 mM, NaCl 1 M) supplied with complete protease inhibitors (Pierce). Cells were lysed by sonication and soluble fraction loaded on Ni Sepharose High Performance resin (Cytiva). After several washes using buffer B (Tris-HCl 20 mM pH 8 TCEP 1 mM, NaCl 1 M, Imidazole 20 mM), elution was performed in buffer C (Tris-HCl 20 mM pH 8 TCEP 1 mM, NaCl 1 M, Imidazole 300 mM). Recombinant protein concentrations were measured using a NanoDrop-2000 spectrophotometer (Thermo Fisher Scientific Inc., Waltham, MA, www.thermofisher.com/).

**Table S1:**
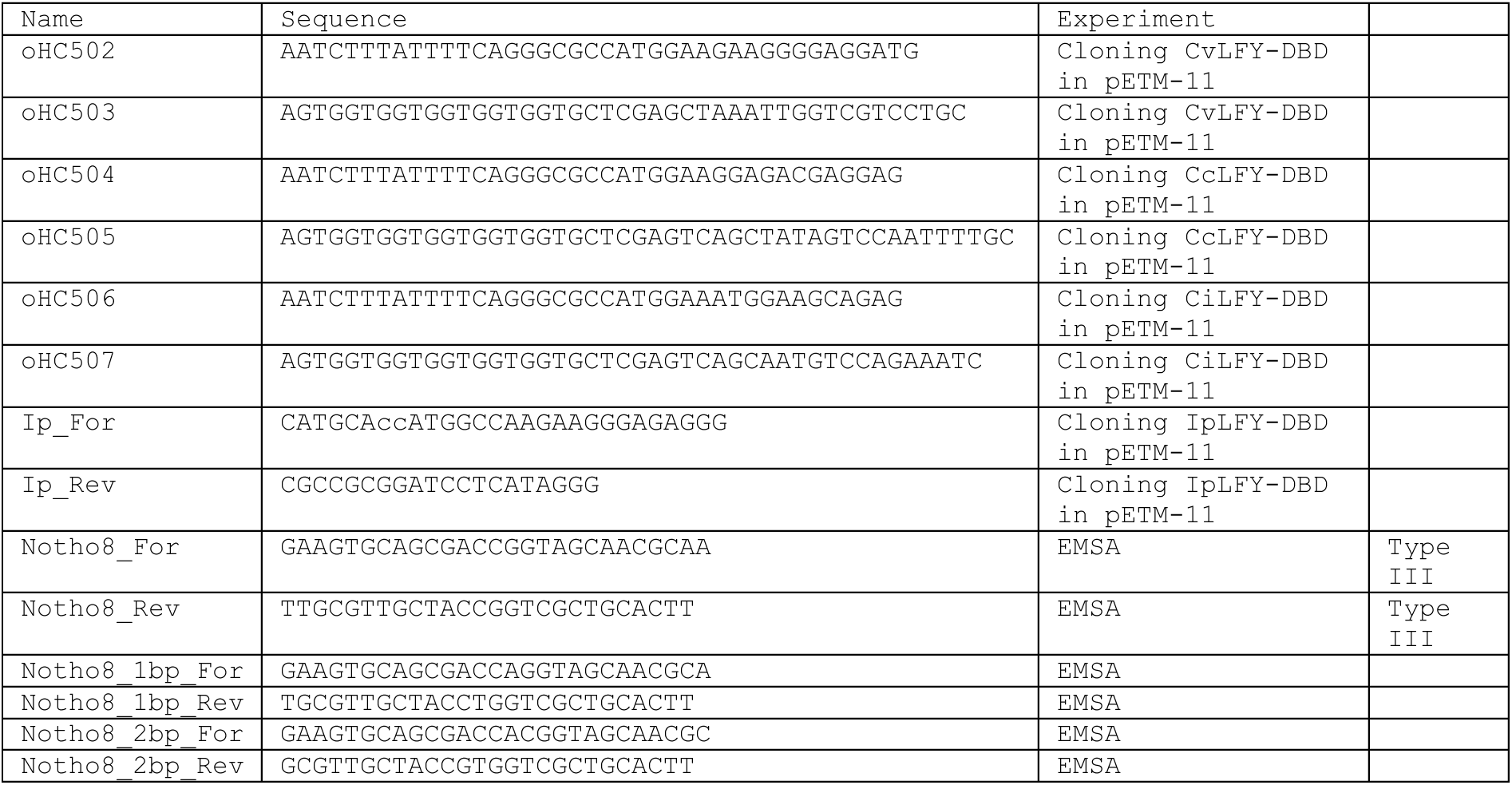
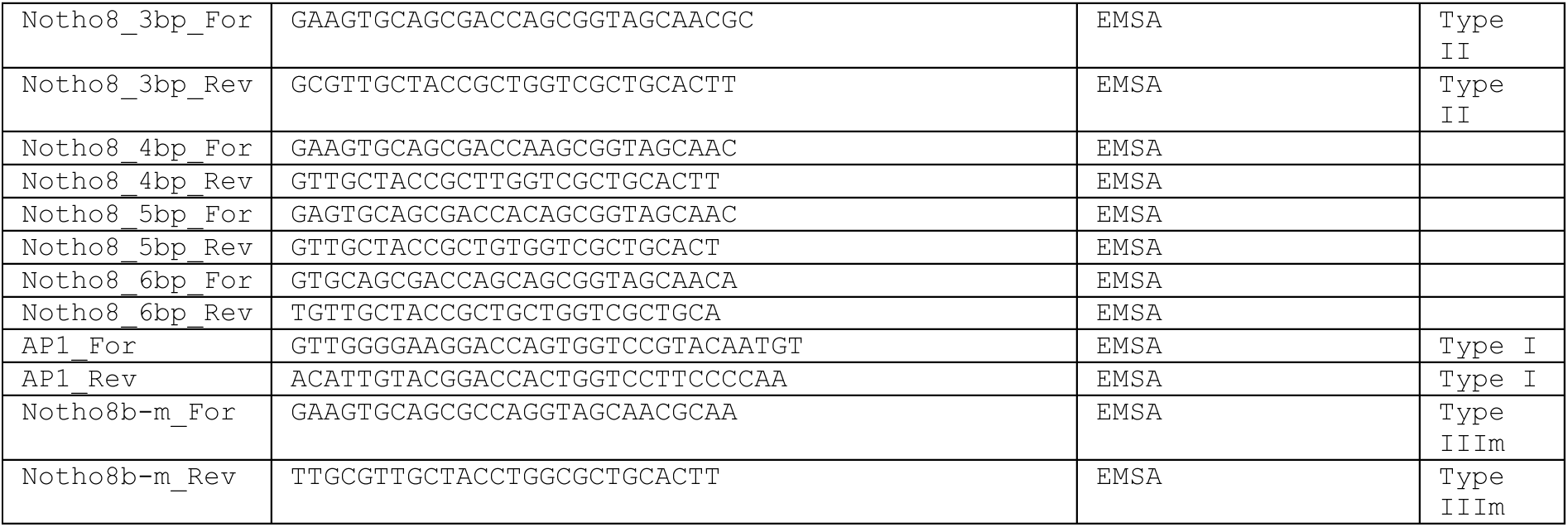
Oligonucleotide sequences.

